# Cross-subunit Interactions that Stabilize Open States Mediate Gating in NMDA Receptors

**DOI:** 10.1101/2020.06.08.140525

**Authors:** Gary J Iacobucci, Han Wen, Matthew B Helou, Wenjun Zheng, Gabriela K Popescu

## Abstract

NMDA receptors are excitatory channels with critical functions in the physiology of central synapses. Their activation reaction proceeds as a series of kinetically distinguishable, reversible steps, whose structural bases are of current interest. Very likely, the earliest steps in the activation reaction include glutamate binding to and compression of the ligand-binding domain. Later, three short linkers transduce this movement to open the gate by mechanical coupling with transmembrane helices. Here, we used double-mutant cycle analyses to demonstrate that a direct chemical interaction between GluN1-I642 (on M3) and GluN2A-L550 (on L1-M1) stabilizes receptors after they have opened, and therefore represents one of the structural changes that occur late in the activation reaction. This native interaction extends the current decay, and its absence predicts deficits in charge transfer by GluN1-I642L, a pathogenic human variant.

**SIGNIFICANCE STATEMENT:** NMDA receptors are glutamatergic channels whose activations control the strength of excitatory synapses in the central nervous system. Agonist binding initiates a complex activation reaction that consists of a stepwise sequence of reversible isomerizations. In addition to previously identified steps in this series, which include agonist-induced closure of the ligand-binding lobes, and the subsequent mechanical pulling by the ligand-binding domain on the gate-forming transmembrane helix, we identify a new cross-subunit interaction, which stabilizes open receptors and slows the rate of the current decay. Naturally occurring NMDA receptor variants lacking this interaction are pathogenic.

## INTRODUCTION

Ionotropic glutamate receptors (iGluRs) are the family of ligand-gated ion-channels responsible for the majority of fast excitatory currents in the central nervous system (1). Among iGluRs, mammalian NMDA receptors are remarkable for their prolonged activations and Ca^2+^-rich currents, biophysical properties that make them directly responsible for synaptic processes that underlie learning, memory and cognition (2). Their biophysical properties vary across development, brain regions, and during physiological and pathological states, due to differential expression of subunits and complex regulatory mechanisms (3). Recently identified NMDA receptor mutations in patient cohorts begin to provide missing links between the physicochemical properties of this critical protein and the physiological and behavioral phenomena they control (4, 5).

NMDA receptors are large heterotetrameric trans-membrane proteins that assemble from two glycine-binding GluN1 subunits, and two glutamate-binding GluN2 subunits. The obligatory GluN1 subunit is practically omnipresent across excitatory synapses, whereas GluN2 subunits, which exist as four distinct subtypes (A-D), have specific expression patterns. Studies with recombinant receptors have demonstrated unique functional features and biological significance for NMDA receptors containing GluN2A or GluN2B subunits (6). Notably, an abrupt replacement of GluN2B subunits with GluN2A subunits marks a critical stage in synaptic development (7). Functionally, GluN2A- and GluN2B-containing receptors have similar pore conductance and calcium permeability properties (8); however, they differ markedly in the time course of their synaptic response, which is fully explained by the distinct kinetics of their activation mechanisms (9). Typically, GluN1/GluN2A receptors produce more robust signals, whereas GluN1/GluN2B receptors are more amenable to structural studies.

All iGluR subunits have modular architectures with large extracellular and cytoplasmic domains and a short, transmembrane domain (TMD), which consists of three transmembrane helices (M1, M3, and M4) and an internally facing P-loop (M2). The external portion of each subunit consists of two stacked globular domains, the N-terminal (NTD) and ligand-binding (LBD) domains, for which several atomic-resolution structures exist (10, 11). In contrast, the cytoplasmic C-terminal domain (CTD), which is least conserved across subunits, appears largely disordered. Although critically important for the receptor’s cellular functions, the NTD and the CTD are dispensable for glutamate-dependent electric activity (12, 13). Therefore, the LBD and TMD layers, and the short linkers that connect them, L1-M1, M3-L2, and L2-M4, perform the essential activation reaction, while the attached NTD and CTD layers provide critical modulation. Consistent with view, the existing evidence suggests that the activation reaction starts with the binding of agonists to the LBDs of resting receptors, which facilitate subsequent local interactions and stabilize LBDs in closed, more compact, and less flexible conformations (2). This whole-domain conformational change repositions the attached LBD-TMD linkers. Notably, the length and mobility of the LBD-TMD linkers correlates with channel opening probability, suggesting they serve as mechanical transducers between LBDs and the TMD-situated gate (14–16). Aside from length and mobility, given their high degree of conservation across species, low toleration to mutations in humans, and structural proximity to the gate, linker residue very likely have additional and more specific roles in receptor activation. Notably, alanine-scanning mutagenesis demonstrate distinct contributions by side-chains of several L1-M1 residues in gating (17, 18). However, because the available structures have limited resolution for these flexible linkers (10, 11, 19–22) the mechanisms by which these side-chains control the gating equilibrium remain unknown.

In keeping with the multiple and diverse intramolecular motions that transform resting receptors into open, current-passing proteins, NMDA receptors have similarly complex kinetic signatures. Statistically derived models, from both single-molecule patch-clamp recordings (23–25) and single-molecule FRET studies (26, 27) have demonstrated that these proteins explore complex free-energy landscapes by transitioning stochastically between several closed and open states, which are distinguishable by their lifetimes. These functionally defined states likely represent entire families of closely related, rapidly interconverting structural conformers. Importantly, these kinetically defined microscopic states occur in a predictable sequence. A current aspiration in this field is to identify the series of structural changes that underlie the functionally observed sequence of lifetime changes, by integrating structural and functional information about receptor activation.

Here we used structure-guided mutagenesis, kinetic modeling of single receptor currents, and double-mutant thermodynamic analyses to show that show that during gating, L550, on GluN2 L1-M1 linker, couples energetically with I642, on GluN1-M3 helix, and that this cross-subunit interaction, which occurs late in the gating reaction, serves specifically to stabilize an open, ion conductive conformation. Our results identify a new cross-subunit state-specific interaction that mediates NMDA receptor gating, and provide insight into how mutations at this site may produce neuropsychiatric pathologies.

## METHODS

### Molecular modeling and targeted MD simulations

We built a closed state model of the NTD- and CTD-lacking GluN1/GluN2A receptor (GluN1 397-838; GluN2A 397-842) with SWISS-MODEL (28–31), using the CTD-lacking GluN1/GluN2B crystal structure (PDB ID 4tlm) from OMP database (32) as template (accession codes P35439, Q00959 for GluN1, GluN2A respectively). We embedded this structure using the Membrane Builder function (28, 33, 34) of the CHARMM-GUI webserver (35, 36) in a bilayer of 1-palmitoyl-2-oleoyl phosphatidylcholine (POPC) lipids surrounded by a box of water, K^+^ and Cl^−^ (0.15 M),with 15Å buffer of water/lipids extending from the protein in each direction. The protein was properly rotated to have a minimal water box size and ~256,000 atoms in total. After energy minimization, we performed six steps of equilibration, which gradually reduced harmonic restraints applied to protein, lipids, water, and ions. Finally, we conducted molecular dynamics (MD) runs in the NPT (isothermal-isobaric) ensemble using the Nosé-Hoover method (37, 38) at 303K. For pressure coupling we used the Parrinello–Rahman method (39); for non-bonded interactions we used a 10-Å switching distance and a 12-Å cutoff distance. For electrostatics calculations we used the particle mesh Ewald method (40). We constrained the hydrogen-containing bond lengths with the LINCS algorithm (41), which allowed a 2-fs time step for the MD simulation. The energy minimization and MD simulation were performed with the GROMACS program (42) version 5.0.3-gpu using the CHARMM36 force field (43, 44) and TIP3P water model (45). To sample the closed state, we conducted three 200-ns MD simulations.

To model a putative open state, we used a modeled ‘active’ conformation of LBD (GluN1 residues 397-441, 451-542, 663-798; GluN2A residues 404-438, 453-537, 659-800) as a template for targeted MD simulations (46) with NAMD V2.9b (47). To model the target LBD conformation, we performed molecular dynamics flexible fitting (MDFF) (48) to flexibly fit an ‘active’ LBD structure (PDB ID: 5FXG) (20) into an active-state cryo-EM map (EMDB ID: 3352). Due to the relatively low resolution of the templates, it was necessary to refine the active-state model of LBDs with MDFF, especially the region of LBD-TMD linkers, which is critically engaged in channel activation. MDFF improved the overall cross-correlation coefficient (CCC) from 0.66 to 0.8. Next, we used the MDFF-refined LBD as target and the closed-state model (see above) as the initial conformation to carry out a 10-ns targeted MD simulation using the Colvars module of NAMD. We applied 200 kcal/mol/deg dihedral restraints to GluN2A M3 helices to ensure the last helical turn is maintained during simulation (49). After the targeted MD simulation, we used the resulting active-state conformation to rebuild the wild type and mutant GluN1/GluN2A with CHARMM-GUI webserver. Finally, we conducted three 200-ns MD simulations using the same MD settings as described above, along with 12 kcal/mol/nm^2^ positional restraints applied on the LBD. In all MD trajectories, only the last 150 ns was kept for energetic analysis and the last 10 ns was used for HOLE calculation (50). To measure inter-residue Van der Waals energies, we used the NAMDEnergy module in the VMD program (47, 51).

### Residue conservation and genetic variation analyses

We used several annotated databases (Uniprot, Ensembl Metazoa) to identify iGluR sequences homologous to human GluN1-1a (Q05586) primary sequence (supplementary table 2). We excluded sequences lacking iGluR-like topology as predicted by TMHMM v2.0 software (http://www.cbs.dtu.dk/services/TMHMM/). In total, we collected 266 iGluR sequences from a diverse set of organisms across the animal, plant and prokaryotic kingdoms spanning an estimated evolutionary divergence time from *homo sapiens* of 4.3 billion years (timetree.org; see supplementary Table 1). We aligned sequences with Clustal Omega v1.2.2 (http://www.clustal.org/omega/), and assigned evolutionary conservation scores to each residue based on conservation of physico-chemical properties using the AMAS method (52) in Jalview v2.10.5 (http://www.jalview.org/).

We examined genetic variation in the human *GRIN2A* gene using the Exome Aggregation Consortium (ExAC) database (v0.3.1), which compiles genomic data from 60,706 unrelated individuals. We restricted our search to variants that passed ExAC quality thresholds. Further, we used these data to calculate missense tolerance ratio (MTR) scores using a sliding window in MTR Viewer (accessed April 2019; http://mtr-viewer.mdhs.unimelb.edu.au/mtr-viewer) (53). Onto this sequence, we superimposed disease-associated mutations annotated in ClinVar (accessed March 2020), as ‘pathogenic,’ ‘likely pathogenic,’ and ‘likely pathogenic/pathogenic.’ A literature search provided additional disease-associated variants.

### Cell culture and molecular biology

HEK-293 cells (CRL-1573, American Type Culture Collection) were grown in DMEM (Invitrogen, Grand Island, NY) with 10% FBS, 1% penicillin-streptomycin, and 10 mM MgCl_2_ and were sustained in 5% CO_2_ atmosphere at 37°C. At ~50% confluence, cells were transfected with rat GluN1-1a (U08261), GluN2A (M91561), and GFP in 1:1:1 ratio using polyethyleneimine (54) (Polysciences, Inc.; 23966-1; lot: 705149). We verified each construct by sequencing before and after subcloning, and after plasmid amplification (QIAGEN, Valencia, CA). All constructs were in pcDNA3.1(+) under control of the CMV promoter. Cells were used 24 - 48 hours post transfection for electrophysiological measurement.

### One-channel current recordings and analyses

We recorded currents from one NMDA receptor with the cell-attached patch clamp technique (55–57). During recordings, cells were bathed in 1X PBS. Recording pipettes (borosilicate, 15–25 MΩ) contained (in mM) 150 NaCl, 2.5 KCl, 10 HEPBS, 1 EDTA, 0.1 glycine, and 1 glutamate, at pH 8 (NaOH). Positive voltage (+100 mV) was applied through the recording pipette (recording potential, ~ −120 mV). Currents were amplified, low-pass filtered (10 kHz; Axopatch 200B) and sampled at 40 kHz (Molecular Devices; DigiData 1440A). All data were acquired with QuB software (University at Buffalo, Buffalo, NY) into digital files and processed off-line.

Data obtained from cell-attached patches containing a single active receptor were minimally processed and idealized, and subjected to kinetic analyses to extract kinetic parameters (58). Data recording, manipulation and analyses were done in QuB. Briefly, recording artifacts, including noise spikes and baseline drift, were eliminated by replacing the sampled amplitudes within spikes with amplitude levels from samples immediately adjacent (open or closed). Baseline drift was corrected by imposing the amplitude level for closed events to baseline at user-defined nodal points as needed. Currents were idealized with the segmental-k-means (SKM) algorithm on digitally filtered data (12 kHz) with no imposed dead time (59). This procedure assigns each sampled data point to either the open (O) or closed (C) class producing a vector of open and closed interval durations, while preserving the temporal sequence in which these events occurred. Kinetic parameters were optimized in QuB by fitting user-defined state models to the sequence of SKM-idealized intervals. Fitting was done by a maximum interval likelihood (MIL) method using an imposed dead time of 75 μs (60). Models were ranked according to their calculated maximum value of the log likelihood (LL) function (23). Bursts were isolated by defining for each recording a critical time (t_crit_) that separates the three shortest closed components from the two longest-lived components, corresponding to desensitized intervals. Kinetic models were fitted to the bursts of activity collected across all recordings obtained for a specific condition. The rates constants resulting from the fitting procedures were used to calculate free energy landscapes according to the relationship ΔG^0^ = −R·T·ln(K_eq_), where R is the molar gas constant, T is the absolute temperature, and K_eq_ is the equilibrium constant for the transition considered. Barrier heights were estimated as E^†^ = ΔG^0^ + (10 – ln(*k*_f_)). Energy differences were calculated relative to state C_3_.

### Double-mutant cycle analyses

To evaluate the extent of energetic coupling between specified residues in wild-type receptors, we built upon the theoretical groundwork put forth by Chowdhury et al. (61). Briefly, receptors were considered to exist in an initial closed state, S_1_, which undergoes n conformational changes through intermediate pre-open states, S_i_, to reach the final open form, S_n+1_. In the context of this system, perturbation of a residue at position A in either the wild-type background or a background that contains a mutation at position B, may result in a change of free energy at any of the intermediates, S_i_, in the activation pathway. Central to the interpretation of mutant cycle analyses is the assumption that for energetically independent positions the observed perturbation energies are equal and additive. Conversely, unequal perturbation energies are indicative of energetic coupling between the two positions. Thus, the energy of interaction at any of the intermediates can be described by:

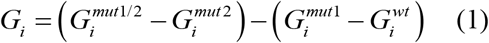

which can be empirically measured by:

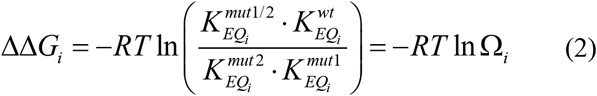

where *K*_*EQ,i*_ refers to the equilibrium constant of the indicated construct and *Ω*_*i*_ refers to the coupling factor of the *i*^*t*h^ transition of the indicated wild-type or mutant construct. The non-additivity of ΔΔG_i_ was determined by:

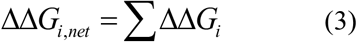

To infer the contribution of interaction energy in stabilizing the transition state between intermediate states in the activation pathway, we estimated the change in activation energy between each state in the forward direction according to:

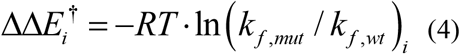

where *k*_*f*_ corresponds to the forward rate constant of transition step *i*. The same analysis was applied in the reverse direction by substituting *k*_*f*_ for the reverse rate constant, *k*_*r*_, of transition step *i*. The non-additivity of the ΔΔE_i_^†^ was then calculated by:

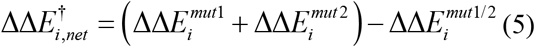

### Simulations of macroscopic current responses

To determine the relative contributions to the overall macroscopic current output for the calculated interaction energies at each transition step, we simulated the macroscopic response of glycine-bound receptors to 1 ms pulse of saturating 1 mM glutamate. All simulations were performed in MATLAB 2017a (Mathworks) using the built-in matrix exponential function, *expm*. We used the rate constants derived in QuB to construct a matrix (*A*) of *n* × *n* size where *n* is the number of states in the model and each element is the rate constant value between the corresponding states. Two glutamate binding steps were appended to C_3_ using experimental binding rate constants derived previously (24). A deterministic simulation of the occupancy of all states with time resolution, *dt*, was performed by iteratively solving:

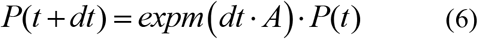

We calculated the final macroscopic current amplitude by summing the occupancies of both open states in the model at each time point. The total charge transferred (*Q*) during the simulation was calculated as the integral over the simulation time using the built-in *trapz* function in MATLAB using the measured unitary Na^+^ conductance for each construct (**Table 1**).

**Table 1:** Kinetic parameters in bursts.

| Receptor | Unitary Current (pA) | Burst P<sub>o</sub> | MOT (ms) | Burst MCT (ms) | Events | N | Total burst time (min) |
| --- | --- | --- | --- | --- | --- | --- | --- |
| GluN1/GluN2A | 8.9 ± 0.6 | 0.73 ± 0.06 | 4.4 ± 0.6 | 1.2 ± 0.6 | 4,663,279 | 8 | 419 |
| GluN1<sup>I642L</sup>/GluN2A | 7.8 ± 0.9 | 0.08 ± 0.03* | 1.3 ± 0.3* | 23.0 ± 4.0* | 118,494 | 7 | 31 |
| GluN1/GluN2A<sup>L550I</sup> | 8.2 ± 0.3 | 0.08 ± 0.03* | 1.7 ± 0.4* | 39.9 ± 10.3* | 22,141 | 7 | 14 |
| GluN1<sup>I642L</sup>/GluN2A<sup>L550I</sup> | 5.4 ± 0.3* | 0.003 ± 0.001* | 0.7 ± 0.1* | 244.8 ± 55.4* | 5,682 | 7 | 21 |
| GluN1<sup>I642A</sup>/GluN2A | 4.4 ± 0.7* | 0.07 ± 0.02* | 1.3 ± 0.2* | 36.4 ± 4.5* | 47,732 | 7 | 17 |
| GluN1/GluN2A<sup>L550A</sup> | 7.0 ± 0.4 | 0.08 ± 0.01* | 1.9 ± 0.1* | 32.4 ± 5.5* | 24,170 | 12 | 10 |
| GluN1<sup>I642A</sup>/GluN2A<sup>L550A</sup> | 5.8 ± 0.3* | 0.033 ± 0.003* | 1.0 ± 0.1* | 1429.3 ± 6.4* | 30,097 | 7 | 12 |

\**p* < 0.05 two-tailed Student *t*-test compared to GluN1/GluN2A
Data represented as mean ± s.e.m

### Statistics

We compared gating parameters (mean open time, mean closed time, open probability, and rate constants) using a paired two-tailed Student’s *t*-test, because they distributed normally as determined by Anderson-Darling test. We displayed results as histograms or using box-and-whisker plots, where whiskers indicate the range, box edges indicate 25^th^ and 75^th^ percentiles, and horizontal red line indicates the distribution mean.

## RESULTS

### Identifying putative state-dependent interactions

Previously, we built a homology model for CTD-lacking GluN1/GluN2A receptors in a closed state and used coarse-grained modeling to envision the pathway of conformational change toward the open state. With this approach we identified a small number of sites where pairs of residues changed their relative positions along the opening trajectory (62). Notably, these pairs clustered spatially, and these hotspots overlapped with patient-derived missense mutations, suggesting that subtle structural perturbations in these regions alter receptor function in biologically significant ways. The low-resolution of the coarse-grained approach we used in this previous work prevented us from observing side-chain motions that we could select for experimental testing. To achieve this goal, we set up to simulate the open/active state with atomic-resolution models. We focused on the smallest receptor assembly that preserves ligand-dependent gating. Such a minimal receptor, GluN1_ΔNΔC_/GluN2A_ΔNΔC_, consists of linked LBD and TMD layers, and lacks the larger NTD and CTD layers (**Figure 1a**, *left*). To reproduce the activation trajectory explored previously with our flexible fitting approach, we targeted the LBD of the resting minimal receptor to a putative ‘active’ conformation (based on PDB: 5FXG) (20). Next, we subjected the system to 200-ns MD simulation with restrained LBDs to relax further the TMD. Comparing initial and final conformations, we observed substantial pore dilation, consistent with progression towards open conformations (**Figure 1a**, *right*). Next, we examined and compared the closed and the open structures to identify pairs of residues that interact specifically in the open state: we keep top 10 pairs with maximal increase in atomic contacts, which are also associated with known disease mutations (supplementary table 1). Among those 10 residue pairs, we selected the GluN1-I642 and GluN2A-L550 pair based on its proximal location to the gate, considerable conservation across species, and potentially pathologic phenotype. The remaining pairs will be investigated in the future.

**Figure 1:**
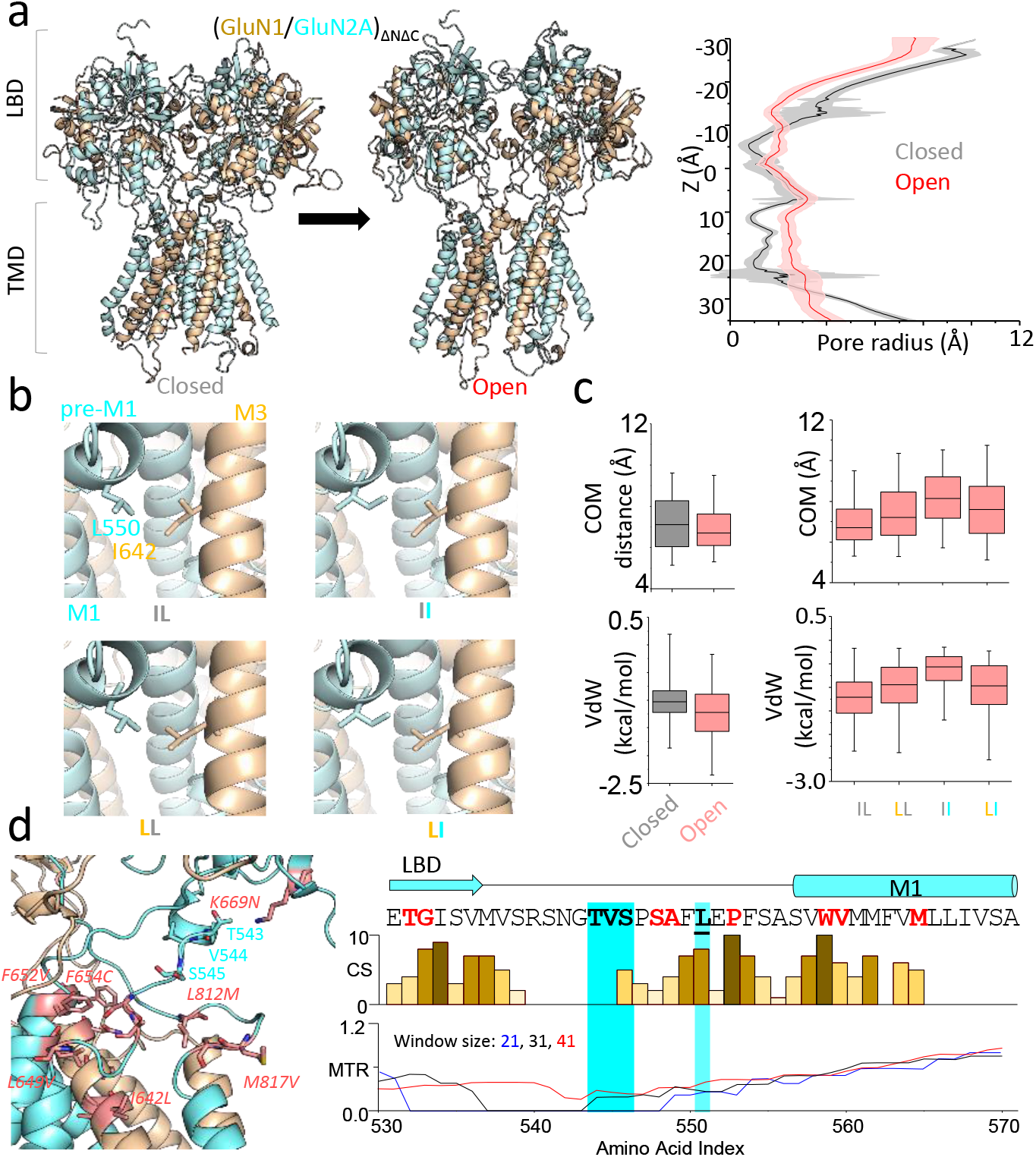
All-atom MD simulation identifies putative state-dependent cross-subunit interaction. (**a**) *Left*, all-atom structural model of (GluN1/GluN2A)_ΔNΔC_ in closed and open conformations. *Right*, HOLE analysis of pore diameter in closed (black) and open (red) conformations (mean ± s.d., 3 runs). **(b)**Close-ups illustrate predicted structural proximity between GluN1-I642 (on LBD/M1-linker, gold) and GluN2A-L550 (on M3-helix, cyan) in the four constructs examined. (**c**) *Left*, measured centers of mass (COM) distances and Van der Waals (VdW) contact energies between GluN1-I642 and GluN2A-L550 sidechains in closed (grey) and open (red) structures. *Right*, conservative substitutions at GluN1-I642 or/and GluN2A-L550 affect COM distance and VdW energy between these residues in the open state, consistent with state-dependent interaction. (**d**) *Left*, GluN2A-L550 maps to a previously identified hotspot (residues labelled in cyan); *right*, hotspot residues are evolutionarily conserved (conservation score, CS), and have low mutation tolerance ratios (MTR, shown with three sliding scales) in human populations. Bold highlights previously identified hotspot residues; red highlights reported and predicted disease-associated mutations

Specifically, the distance between GluN1-I642 and GluN2A-L550, measured as the average side-chain center-of-mass (COM) distance, which was quite short in the closed structure (7.1 ± 1.2 Å) became even shorter in the open structure (6.9 ± 1.1 Å) (**Figure 1b, c**). In this range, the shorter COM distance may reflect or not a change in chemical coupling between the two residues, and this coupling can be attractive or repulsive. The measured average Van der Waals (VdW) contact energies became more negative, changing from −1.02 ± 0.37 kcal/mol in the closed to −1.22 ± 0.50 kcal/mol in the open conformation (**Figure 1c***, left*). Although small, this change in energy is consistent with the formation of favorable contacts between these residues during opening, with a stabilizing effect on the open structure. If indeed, the closer proximity of the two residues in the open state is indicative of a favorable mutual interaction, both the COM distance and the VdW contact energy would depend on the identity of the side chains involved. To probe for this, we repeated the MD simulation after introducing simple side-chain isomerizations at these sites: GluN1-I642L or/and GluN2A-L550I. We found that these conservative substitutions increased the COM distance between these residues in open receptors to 7.4 ± 1.3 Å for GluN1-I642L (LL), and to 8.1 ± 1.2 Å for GluN2A-L550I (II); whereas in the double mutant (LI), which inverted the residues at these two adjacent sites, this distance was 7.6 ± 1.4 (**Figure 1c**, *right*). Similarly, substitutions decreased the negative VdW contact energy in open receptors to −0.99 ± 0.52 kcal/mol for GluN1-I642L (LL), and to −0.63 ± 0.36 kcal/mol for GluN2A-L550I (IL); whereas the double mutant (LI) returned the contact energy to a more wt-like value, −1.03 ± 0.57 kcal/mol; (**Figure 1c**, *right*). These simulations, although limited in scope, are at a minimum internally consistent with a scenario where during activation, GluN1-I642 and GluN2A-L550 move closer to each other, and can form favorable side-chain specific interactions, which may add to the stability of open states.

Further, the location, conservation, and pathogenic potential of these residues support an important role for these residues in receptor function. Both GluN1-I642 and GluN2A-L550 are proximal to the receptor gate. GluN1-I642 resides on the highly conserved M3 transmembrane helix, just one turn below the invariant SYTANL**<u>A</u>**AF sequence, which forms the ligand-controlled pore gate at its +6 position (63). Likewise, GluN2A-L550 maps to the highly conserved L1-M1 linker **(Figure 1d**, *right*), which is critical to gating (17, 18, 64, 65), and hosts many recently identified pathogenic missense mutations (**Figure 1d**, *left*) (17, 62). Lastly, whole genome sequencing in select patient populations identified the GluN1-I642L variation as likely pathogenic (GluN1^I642L^; ClinVar ID: 421935). Together, these considerations provided the motivation to investigate these two side chains and their potential interaction with in-depth functional studies.

### Each GluN1-I642 and GluN2A-L550 contribute to gating in wild-type full-length receptors

To explore whether the subtle structural alterations produced by conservative substitutions at GluN1-I642 and GluN2A-L550 have any functional consequences on the gating of full-length receptors residing in live cells, we set up to record currents produced by substituted receptors and to quantify their reaction mechanisms relative to that of wild-type receptors. We recorded unitary currents from cell-attached patches containing one copy of each full-length wild type GluN1/GluN2A (IL), GluN1^I642L^/GluN2A (LL), or GluN1/GluN2A^L550I^ (II). Kinetic analyses and modeling showed that relative to wild type receptors, each GluN1^I642L^/GluN2A, and GluN1/GluN2A^L550I^ had profoundly changed activity patterns (**Figure 2a**, **Table 1**, **2**). Both mutations caused drastic (~8-fold) decrease in open probability (Po): from 0.7 ± 0.1, for wild type to 0.1 ± 0.0 for I642L (*p* = 7E-4) and L550I (*p* = 5E-3). Mean open durations were ~4-fold shorter, decreasing from 4.5 ± 0.7 ms for wild type to 1.3 ± 0.3 ms (*p* = 8E-4) for GluN1-I642L, and 1.9 ± 0.4 ms (*p* = 2E-3) for GluN2A-L550I. Mean closed durations within bursts were ~10-fold longer, increasing from 1.3 ± 0.3 ms for wild-type to 23.0 ± 3.9 ms for I642L (*p* = 1E-5) and 39.9 ± 10.3 ms for L550I (*p* = 3E-5) (**Figure 2b**, **Table 1**).

**Figure 2:**
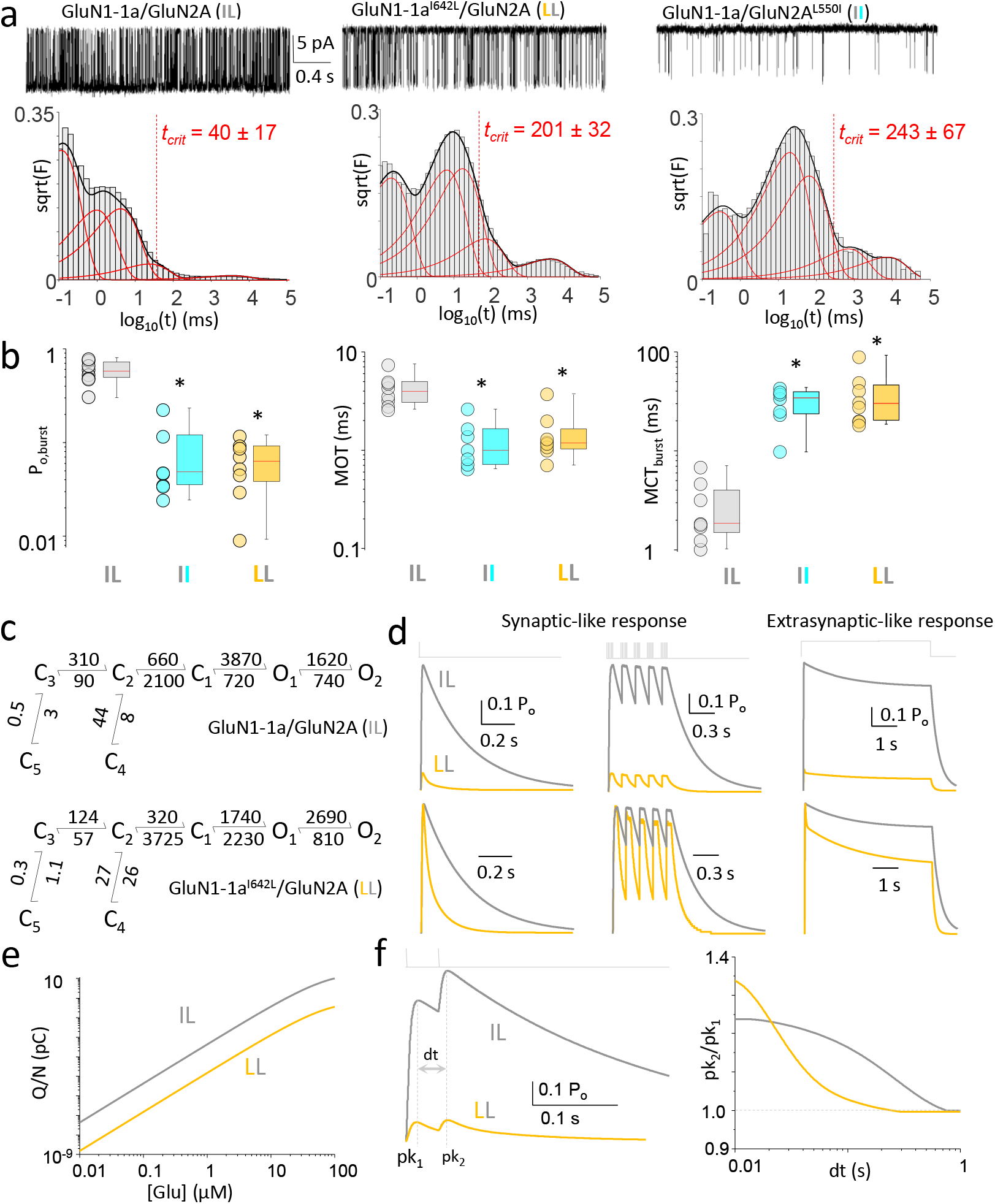
Both GluN1-I642 and GluN2A-L550 contribute to gating in full-length receptors. (**a)** Unitary current recordings (3 s, *top*) and corresponding closed event duration histogram (*below*) for wild-type I642L L550I (IL; *n* = 8), GluN1^I642L^/GluN2A (LL; *n* = 7), and GluN1/GluN2A^L550I^(II; *n* = 8) receptors. Bursts were defined using duration thresholds (t_crit_) calculated for each recording. (**b**) Plots illustrate open probability (Po), mean open (MOT) and mean closed (MCT) times within bursts of activity in each recording. \**p* < 0.05, two-tailed Student’s t-test, relative to wild-type. (**c**) Reaction mechanisms determined from global fits to data obtained from wild-type (IL) and the pathologic GluN1^I642L^-containing receptors (LL). (**d**) Predicted macroscopic responses to synaptic-like (1 ms) or theta-burst stimulation, and to extra-synaptic- like (5 s) stimulation with glutamate (1 mM). (**e**) Predicted total charge transfer (Q_T_/N) per channel across total simulation in response to one synaptic stimulus (1 ms). (**f**) Predicted current response to a two-pulse stimulus (*left*). Fractional current potentiation (Pk_2_/Pk_1_) in response to a two-pulse protocol relative to inter-pulse duration (dt).

**Table 2:** Time components within bursts.

| Receptor | Closed components (ms) |  |  |  | Open components (ms) |  |  |
| --- | --- | --- | --- | --- | --- | --- | --- |
|  | τ<sub>1</sub> | τ<sub>2</sub> | τ<sub>3</sub> | τ<sub>fast</sub> | τ<sub>Low</sub> | τ<sub>Medium</sub> | τ<sub>High</sub> |
| GluN1/GluN2A | 0.1 | 1.3 | 5.3 | 0.1 | 2.6 | 6.3 | 13.8 |
| GluN1<sup>I642L</sup>/GluN2A | 0.2 | 9.1 | 38.5 | 0.2 | 1.6 | 4.2 | -- |
| GluN1/GluN2A<sup>L550I</sup> | 0.3 | 20.1 | 81.4 | 0.5 | 3.4 | -- | -- |
| GluN1<sup>I642L</sup>/GluN2A<sup>L550I</sup> | 0.2 | 138.1 | 525.3 | 0.8 | 6.8 | -- | -- |
| GluN1<sup>I642A</sup>/GluN2A | 0.3 | 7.2 | 27.3 | 1.3 | 3.3 | -- | -- |
| GluN1/GluN2A<sup>L550A</sup> | 0.1 | 11.7 | 46.0 | 0.2 | 2.1 | -- | -- |
| GluN1<sup>I642A</sup>/GluN2A<sup>L550A</sup> | 0.2 | 7.4 | 31.8 | 0.1 | 0.9 | 2.1 | -- |

The GluN2A-L550 position is invariant in human populations, and the conservative GluN1-I642L is pathogenic. The functional data we accumulated GluN1-1a^I642L^/GluN2A allows us to derive its kinetic mechanisms (**Figure 2c**) and predict how this conservative substitution may affect the macroscopic responses. Simulations with several physiologic stimulation patterns show that responses form pathogenic receptors (LL) were drastically smaller in amplitude and decayed substantially faster, relative to wild-type receptors (IL) (**Figure 2d**), resulting in be ~100-fold smaller charge transfer over a wide range of glutamate concentrations (**Figure 2e**), and that they would have altered frequency-dependent potentiation (**Figure 2f**) (2, 24). Cellular consequences of these biophysical changes may include deficits in signal integration across dendritic arbors and altered short-term and long-term synaptic plasticity, with likely repercussions on overall excitability, information processing and storage.

These results demonstrate that the side chains of both GluN1-I642 and GluN2A-L550 are critically involved in prolonging openings and shortening closures and are consistent with the mechanism suggested by MD simulations, in which the interaction between these side chains stabilizes open states. However, these results do not prove direct interaction between the two.

### GluN1-I642 with GluN2A-L550 interaction contributes to gating

To determine whether GluN1-I642 and GluN2A-L550 interact functionally during gating, we set up to perform double-mutant cycle analysis on this residue pair (66). In this classic approach, if the changes in free energy caused by the double mutation is simply the additive effect observed with single mutations, the residues likely contribute independently to the gating reaction. In contrast, if the double mutation causes changes in free energy that are distinct from the sum of changes observed with single mutations, it is likely that the two residues interact during the observed process, and the magnitude of the difference correlates with the strength of the interaction.

For these thermodynamic analyses, we used initially the records obtained above for wild-type GluN1/GluN2A (IL), GluN1^I642L^/GluN2A (LL), and GluN1/GluN2A^L550I^ (II). In addition, we recorded from GluN1^I642L^/GluN2A^L550I^ (LI). To mimic more closely the closed-to-open trajectory simulated by the MD trajectories, we selected from these records, bursts of activity, which represent repeated stochastic excursions between resting and open states. This was done by identifying in each record, and excluding from analyses, periods when the receptors were closed for long time periods, indicative of sojourns into desensitized states *(57, 67, 68)* (**Figure 3**, **Table 1**, **2**). Kinetic modeling of activity within bursts revealed that relative to single mutations, the double mutation produced a larger, synergistic increase in the free energy of gating (1.66 kJ mol^−1^) indicative of strong functional coupling between the queried residues (**Figure 3a**). This value is within reported energies for Van der Waals contacts (0.4 - 4 kJ mol^−1^) and considering our predicted COM distance between these residues (6.9 Å), it is consistent with an attractive force that would stabilize the open state.

**Figure 3:**
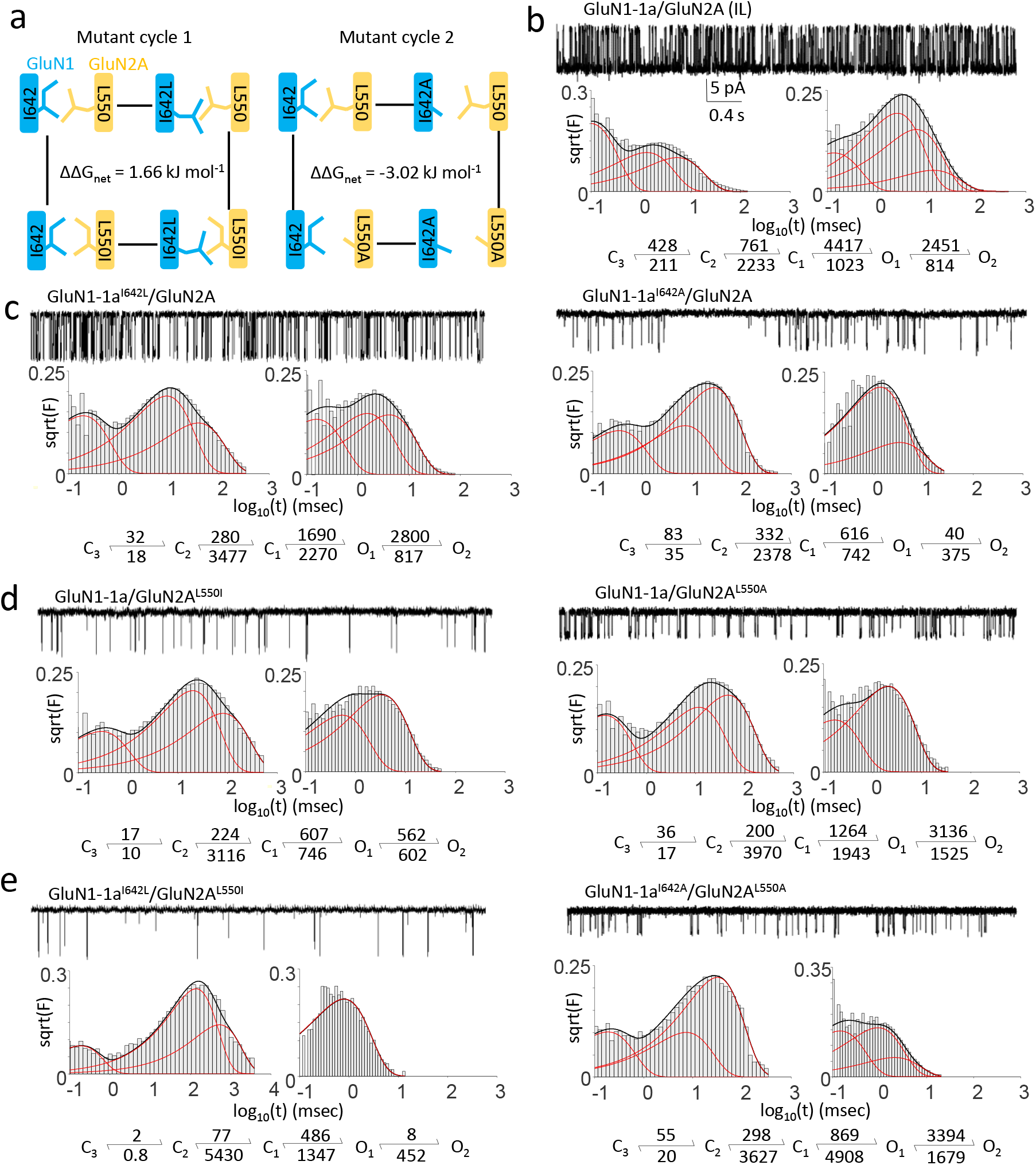
Double-mutant cycle analyses for GluN1^I642^ and GluN2A^L550^. (**a**) Diagrams for the two cycles examined, with calculated free energy changes (ΔΔG_net_) due to mutual interactions during gating (**b - e**). Representative 3-sec periods of continuous single-channel currents, with event duration histograms, and kinetic activation mechanism for the indicated receptors. All rate constants (s^−1^) were derived from global fits of the indicated models to burst-data pooled across all the recordings for each construct.

Classic double-mutant cycle analyses, rather than introducing complementary mutations as we have done above, typically use alanine as the substitution of choice, to eliminate both specific and non-specific putative interactions contributed by the residues under observation. Therefore, we repeated our measurements and analyses with receptors containing GluN^I642A^, GluN2A^L550A^, or the double mutation GluN1^I642A^/GluN2A^L550A^ (**Figure 3**, **Table 1**, **2**). For this more drastic structural change, we calculated that interactions between the side chains of GluN1-I642 and GluN2A-L550 contribute a total of 3.02 kJ mol^−1^ to the gating reaction. Given the observation from MD simulation that simple side-chain isomerizations (leucine/isoleucine mutagenesis) shifted the overall center-of-mass distance and VdW contact energy toward less stable open configurations (**Figure 1d**), we surmise that specific contacts between GluN1-I642 and GluNA2A-L550 contribute a sizeable stabilizing interaction to the open state. Moreover, the larger change observed with side-chain truncations (alanine mutagenesis) suggests that non-specific steric effects by these side chains are also contributing to open sate stability. Based on these results, we suggest that GluN1-I642 and GluN2A-L550 side chains facilitate the NMDA receptor gating reaction substantially, through both their direct specific and nonspecific contacts with each other during gating.

### GluN1-I642 with GluN2A-L550 interaction catalyze opening and stabilize open receptors

NMDA receptors have complex activation reactions whose adequate kinetic description must include at a minimum three closed states and two open states (57). Although the structural correlates of these statistically-defined states are unknown, a multi-state kinetic model accounts well for all observed microscopic and macroscopic receptor behaviors (24). Importantly, it also shows how the channel’s biophysical properties translate into physiological functions at synapses (2). Therefore, in addition to considering a global closed-open gating reaction for each arm of the thermodynamic cycle, as in **Figure 3a**, we aimed to resolve contributions by the interaction of GluN1-I642 with GluN2A-L550 to individual kinetic steps along the activation pathway. We used the rate constants estimated as in **Figure 3b-e**, to calculated the change in free energy (ΔΔG_int_) contributed by single and double mutations, and therefore the coupling constants (θ) between the two residues at each step along the gating sequence (**Figure 4a**). With this more detailed analysis, we found that perturbing the side chains of GluN1-I642 and GluN2A-L550 affected all the transitions considered. However, the final step (O_1_⇄O_2_) was most drastically changed. During this step, the native interactions stabilized open states, as suggested by MD simulations, and lowered the energy barrier for the O_1_⇄O_2_ transition, indicating a catalytic component for this contribution.

**Figure 4:**
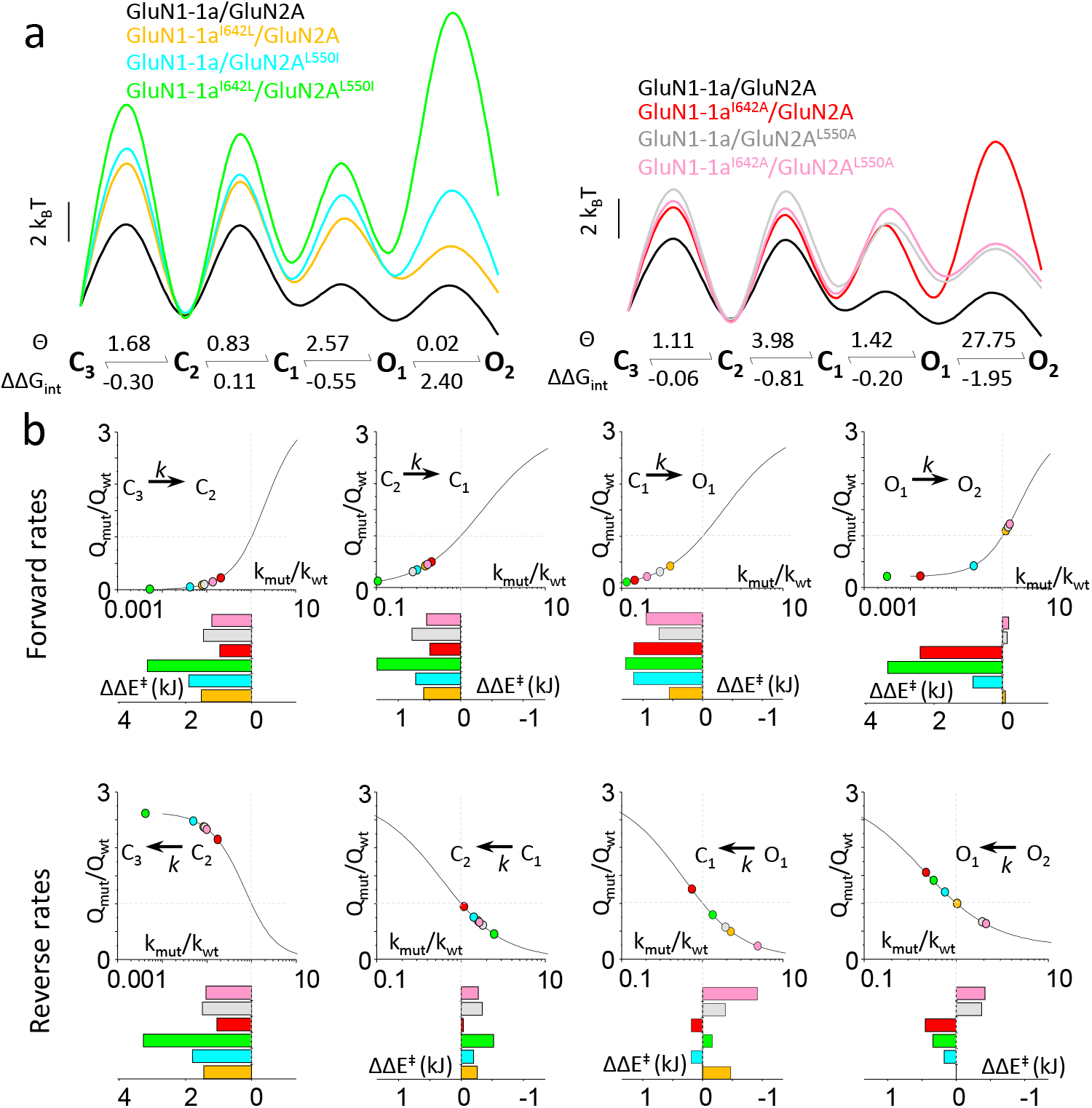
Interactions between GluN1-I642 and GluN2A-L550 stabilize open states and decrease barriers to opening. (**a**) Effects of side-chain isomerization (*left*) and side-chain truncation (*right*) on free energy landscapes (*top*) calculated from measured rate constants (given in **Figure 3**), and break-down of coupling constants (Θ) and free energy contributions (ΔΔG_int_ in kcal/mol) per each transition (*below*). (**b**) In each panel, the indicated wild-type rate (k_wt_) was scaled iteratively by an order of magnitude (k_mut_) and the resulting kinetic model was used to calculate the change in charge transfer (Q_mut_/Q_wt_) following synaptic-like stimulation. Lines represent exponential fits to the simulated data points, and experimentally measured rate constants are superimposed as colored circles. Bar graphs below illustrate the change in the height of the transition state energy ΔΔE^‡^(kcal/mol) for each mutant.

**Figure 5:**
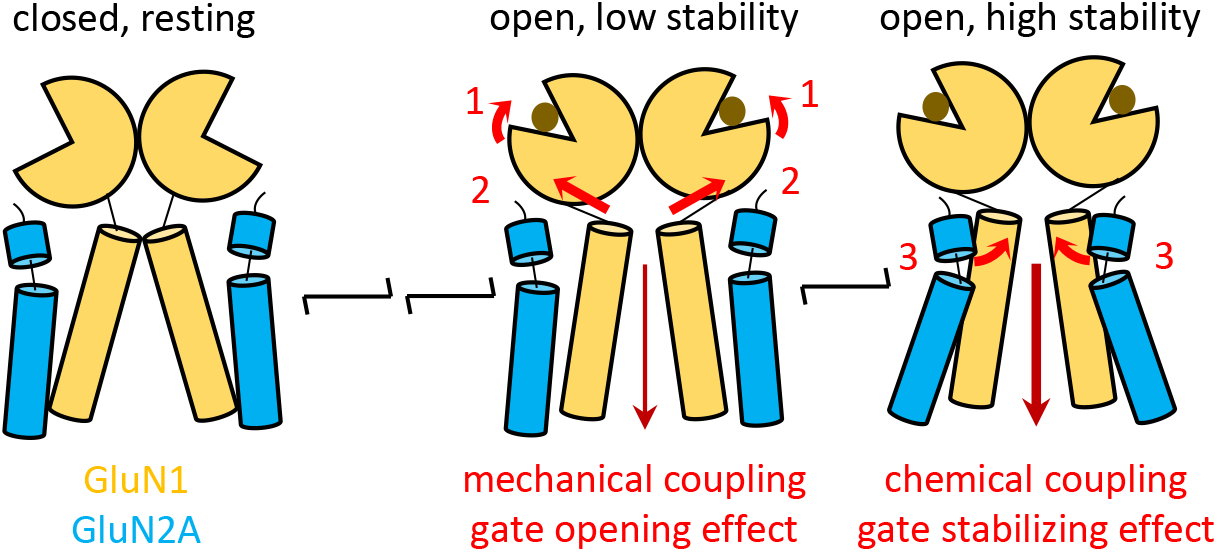
Differential roles of linkers during gating. As receptors transition into open states upon agonist binding and cleft closure (1), mechanical tension on M3 linkers transmits the energy of ligand binding to M3 helices to open the gate (2). With the gate open, residues on M3 below the gate become more accessible to residues on the pre-M1 helix of the GluN2A subunit belonging to the opposing heterodimer, thus stabilizing the gate in an open position (3).

In this study, we focused specifically on contributions by GluN1-I642 and GluN2A-L550 side chains to the gating reaction. Therefore, we recorded currents in divalent-free external solutions, which bypass complexities to the single-channel record contributed by possible changes in calcium permeability, calcium-dependent block and inactivation, or voltage-dependent block by magnesium. However, we noted that doubly substituted receptors displayed unitary currents with amplitudes consistently lower relative to wild type channels, suggesting possible contributions by these residues to the physicochemical properties of the permeation pathway (**Table 1**). This is consistent with previous work suggesting pharmacological modulating in this region can alter permeation (69). Of the singly substituted constructs tested, GluN1-I642A was sufficient to produce the same reduction in current amplitude as the double alanine substitution, suggesting that GluN1 residues on M3 are dominant in setting channel permeability, consistent with previous observations from receptors with Lurcher-type mutations (70).

### State-specific interaction differentially stabilize activation and deactivation transitions

In addition to altering gating kinetics by destabilizing active conformations as expected, disrupting the native interactions between GluN1-I642 and GluN2A-L550 also increased transition barriers along the entire gating reaction, and affected asymmetrically the forward and reverse pathway. This observation has implications for understanding the subtle internal dynamics that make this receptor’s gating sequence, and therefore the mechanistic basis of their macroscopic output, which drives synaptic physiology. A theoretical study examining the effect of state-dependent perturbations at NMDA receptors found that kinetic changes of similar magnitude affect differentially the receptor’s electrical output according to the specific kinetic transition affected (71). Therefore, we set up to estimate how the observed changes in each rate constant, and the corresponding change in activation energy, individually impacted the resulting the magnitude of the macroscopic response.

First, we used the rate constants measured for wild-type receptors (**Figure 3b**) to simulate macroscopic currents in response to synaptic-like stimulation and calculated the resulting charge transfer (*Q_wt_*), as one measure of potential impact on synaptic physiology (2). Next, we scaled each rate constant 10-fold in both directions to estimate the sensitivity of charge transfer (*Q_n_/Q_w_*) to changes in specific rate constants (**Figure 4b**, black lines). Onto the estimated charge vs. rate relationship, we mapped the empirical rate constants measured for the six mutants examined in this study. This analysis shows that absent the native interactions afforded by GluN1-I642 and GluN2A-L550, all forward rates were substantially slower, indicative of larger activation barriers, and resulted in substantial decrease in charge transfer. Both side-chain isomerization (I/L) and side-chain truncations (I/A, L/A) were influential, suggesting that both specific and non-specific interactions at this sensitive site contribute to channel activation in wild-type receptors. In contrast, changes in the reverse (deactivation) rates were generally smaller and had only mild influence on charge transfer, except for the last step in the deactivation sequence (C_3_ ← C_2_), for which the rate was substantially slower. This last step, represents functionally a transition from receptors with very high glutamate affinity (closed LBD lobes), into receptors from which glutamate can dissociate with measurable rates (60 s^−1^/site) (open LBD lobes) (23, 24); it sets the kinetics of macroscopic decay, and for this reason it has steep influence on charge transfer. Together, these simulations point to the possibility that the interactions examined make asymmetric contributions to the opening and closing reactions. This possibility merits further investigation. Nevertheless, the pattern of influence on specific rate constants is consistent with the hypothesis suggested by our initial all-atom MD simulation that GluN1-I642 and GluN2A-L550 form specific state-dependent interactions that stabilize open states and shift the closed-open gating equilibrium to support longer responses.

## DISCUSSION

In this study, we used structure-guided mutagenesis, kinetic modeling, and thermodynamic analyses to identify the first, to our knowledge, open state-specific interaction in NMDA receptors and characterize its mechanism and energetic contributions to gating. We selected this residue pair for in-depth functional studies based on our preliminary results from targeted MD simulations, and their critical proximity to the channel gate, stringent conservation across species, and mutation intolerance in humans. Briefly, targeted-MD simulations suggested that during gating GluN1-I642 and GluN2A-L550 might move closer to each other and form stabilizing interactions. Both GluN1-I642 and GluN2A-L550 map to regions of primary sequence that are highly conserved in iGluR subunits across species and have low mutation tolerance ratios in human populations (64, 72, 73). Specifically, GluN1-I642 resides deeply within the M3 helix, below the strictly invariant SYTANL**<u>A</u>**AF sequence, which hosts the ligand-controlled gate at the +6 position (70, 73, 74). Notably, based on exome sequencing analyses in large patient populations, the GluN1-I642L variant is likely pathogenic (75). Conversely, GluN2A-L550 resides on the L1-M1 transmembrane helix whose residues are critical for gating in all iGluRs (17, 18, 65). This evidence, although circumstantial, motivated us to investigate the role of this residue pair with functional studies.

Our single-channel current recordings show that the side chains of each GluN1-I642 and GluN2A-L550 serve to increase channel open probability by extending channel openings and shortening channel closures. This observation indicates that each of these residues likely forms state-dependent interactions to tilt the gating landscape towards open states. Further, thermodynamic analyses of two mutant cycles show that these two residues interact with each other, and this mutual interaction makes a substantial energetic contribution to the overall gating reaction. Lastly, modeling of single-channel currents with multi-state kinetic mechanisms shows that this interaction catalyzes a gating transition that occurs late in the gating reaction, and among the many glutamate-activated states, it stabilizes specifically a conductive conformation.

Theoretically, every single interaction that changes during the resting-to-open conformational trajectory of NMDA receptors contributes energetically to gating. Numerous studies have shown that glutamate binding across two mobile lobes of the LBD is very likely the first step in this complex sequence (14, 24, 76, 77). This initial bimolecular event brings the two lobes in proximity such that intermolecular interactions can now form between lobes with higher probability (78–80). The cross-lobe interactions are state-dependent in that they bind to and stabilize the closed-lobe, compact LBD conformation, and in so doing make subsequent gating events more probable. In structural parlance, this closed-lobe conformation is that of an ‘active’ receptor even if the channel gate is not yet open. This notation is correct because the closed-lobe arrangement will remain largely unchanged throughout the remaining steps of the gating cycle, including for receptors with open pores. Given that NMDA receptors have four LBDs, and that full LBD layer occupancy is required for measurable open probability, likely four ligands have to bind and four LBD lobes must close, in an unknown order, before the mechanical tension transmitted through LBD-TMD linkers can pull open the four M3 helices to allow ionic permeation. For this reason, one assumes that lobe closure occurs early during the gating reaction and channel opening occurs later. Functional studies support this temporal sequence and the multi-state gating models we used in this study incorporates these observations. The higher resolution afforded by a multi-state reaction mechanism allowed us to conclude that the I642/L550 interaction we studied here, is not only state-dependent in that it stabilizes an ‘active’ conformation, but it is specifically open-state dependent, because the kinetic state it favors is functionally permeable to ions. To our knowledge, this is the first demonstration of an open-state specific interaction in NMDA receptors.

Notably, this cross-subunit interaction connects GluN1 and GluN2A subunits that belong to separate LBD heteromers, reminiscent of the subunit swap documented between the heterodimers of the LBD and NTD layers (10, 11). A conservative substitution at one of the residues in the pair, GluN1-I642L was previously classified as pathogenic. Our results suggest a mechanism for the pathogenicity of this variant. We show that the I-to-L isomerization disrupts native cross-subunit interactions and that this subtle perturbation causes a dramatic decrease in the stability of open states. At the microscopic level, receptors have drastically reduced open probabilities, shorter openings, and longer closures. At the macroscopic level, GluN1^I642L^/GluN2A receptors produce smaller, faster decaying currents, and consequently mediate severely reduced charge transfer. Consistent with these predictions, mutations at the partner residue in this pair (L550A) also produces faster decaying currents, suggesting similar mechanism (17, 18). Given the importance of these biophysical properties of the NMDA receptor response for the receptor’s physiologic role, and given the ubiquitous expression of GluN1 subunits in the central nervous system, it is plausible that the GluN1-I642L variant will cause notable changes in excitability, synaptic integration and plasticity, and information processing.

Overall, our results provide direct empirical evidence for a new mechanism by which side chains of TMD-LBD linker residues contribute to NMDA receptor gating. In addition to the direct mechanical coupling provided by L2-M3 linkers between same-subunit LBD-TMD (14), we demonstrate here a catalytic mechanism by which residues on the L1-M1 linkers connect directly with M3 residues across subunits, and across heterodimers. Therefore, linkers play multiple roles in the gating sequence by providing both necessary mechanical tension to open the gate and later, chemical coupling with residues below the gate to stabilize the open-gate conformation. Specifically, our results show that the interaction between GluN1-I642 and GluN2A-L550 provides a cross-subunit cross-dimer interaction that is intrinsic to the activation sequence and contributes to the characteristically long durations of NMDA receptor activations.

A remaining gap in understanding NMDA receptor gating is the tenuous correspondence between experimentally defined structural conformations and functionally defined kinetics states. Advances in computational power, accumulating structural information, and high resolution kinetic analyses promise to integrate structural and functional views to generate a congruent picture for this receptor’s mechanism. Combining this mechanistic knowledge with clinical observations of patients carrying pathologic variants will bring to fruition the translational potential of this research.

## ACKNOWLEDGEMENTS

We thank Dr. Jamie Abbott for helpful discussion and critiques of the manuscript. We thank Evan Synor and Michael Steward for contributions to data collection. The NIH supported this project through awards R01NS097016 and R01NS108750. G.J.I., H.W., W.Z., and G.K.P conceived the project and designed experiments; G.J.I., M.H. and H.W. collected data and analyzed results. G.J.I. and G.K.P prepared the manuscript. The authors declare no conflicts of interest.

## Notes

### Competing Interest Statement

The authors have declared no competing interest.

