## Supplemental Tables for "Cross-subunit Interactions that Stabilize Open States Mediate Gating in NMDA Receptors"

### Supplementary tables

*Table S1: A list of ten MD-observed residue pairs specific to the open state*

| Pair | Partner A | Partner B | Disease Variant |
|---|---|---|---|
| 1 | GluN1 I642 | GluN2A L550 | GluN1$^{I642L}$ |
| 2 | GluN1 T651 | GluN2A L812 | GluN2A$^{L812M}$ |
| 3 | GluN1 I565 | GluN2A L812 | GluN2A$^{L812M}$ |
| 4 | GluN1 V635 | GluN2A L824 | GluN2B$^{L825V}$ |
| 5 | GluN1 N803 | GluN2A P552 | GluN2A$^{P552R}$ |
| 6 | GluN1 V816 | GluN2A V557 | GluN2B$^{V558I}$ |
| 7 | GluN1 L562 | GluN1 Y647 | GluN1$^{Y647S}$ |
| 8 | GluN1 W563 | GluN1 Y647 | GluN1$^{Y647S}$ |
| 9 | GluN2A P552 | GluN2A F652 | GluN2A$^{P552R}$ and GluN2A$^{F652V}$ |
| 10 | GluN2A P552 | GluN2A N648 | GluN2A$^{P552R}$ |

Table S2: iGluR amino acid sequences used in conservation analysis

| Gene name | Accession number | Database | Species |
|---|---|---|---|
| GRIN1-1a | XM_005266073.4 | https://www.ncbi.nlm.nih.gov/protein/ | *Homo sapiens* |
| GRIN2A | NP_000824.1 | https://www.ncbi.nlm.nih.gov/protein/ | *Homo sapiens* |
| GRIN2B | NP_000825.2 | https://www.ncbi.nlm.nih.gov/protein/ | *Homo sapiens* |
| GRIN2C | AAI40802.1 | https://www.ncbi.nlm.nih.gov/protein/ | *Homo sapiens* |
| GRIN2D | NP_000827.2 | https://www.ncbi.nlm.nih.gov/protein/ | *Homo sapiens* |
| GRIN3A | NP_597702.2 | https://www.ncbi.nlm.nih.gov/protein/ | *Homo sapiens* |
| GRIN3B | NP_619635.1 | https://www.ncbi.nlm.nih.gov/protein/ | *Homo sapiens* |
| GRIA1 | NP_000818.2 | https://www.ncbi.nlm.nih.gov/protein/ | *Homo sapiens* |
| GRIA2 | NP_000817.2 | https://www.ncbi.nlm.nih.gov/protein/ | *Homo sapiens* |
| GRIA3 | AAA67923.1 | https://www.ncbi.nlm.nih.gov/protein/ | *Homo sapiens* |
| GRIA4 | NP_000820.3 | https://www.ncbi.nlm.nih.gov/protein/ | *Homo sapiens* |
| GRIK1 | NP_000821.1 | https://www.ncbi.nlm.nih.gov/protein/ | *Homo sapiens* |
| GRIK2 | NP_001159719.1 | https://www.ncbi.nlm.nih.gov/protein/ | *Homo sapiens* |
| GRIK3 | NP_000822.2 | https://www.ncbi.nlm.nih.gov/protein/ | *Homo sapiens* |
| GRIK4 | NP_055434.2 | https://www.ncbi.nlm.nih.gov/protein/ | *Homo sapiens* |
| GRIK5 | NP_002079.3 | https://www.ncbi.nlm.nih.gov/protein/ | *Homo sapiens* |
| GRID1 | NP_060021.1 | https://www.ncbi.nlm.nih.gov/protein/ | *Homo sapiens* |
| GRID2 | AAH99652.1 | https://www.ncbi.nlm.nih.gov/protein/ | *Homo sapiens* |
| 1281696 | XP_321646.3 | https://www.ncbi.nlm.nih.gov/protein/ | *Anopheles gambiae* |
| 1275196 | XP_314428.4 | https://www.ncbi.nlm.nih.gov/protein/ | *Anopheles gambiae* |
| 1276211 | XP_315527.3 | https://www.ncbi.nlm.nih.gov/protein/ | *Anopheles gambiae* |
| 1272393 | XP_311343.4 | https://www.ncbi.nlm.nih.gov/protein/ | *Anopheles gambiae* |
| GLURIIb | A0A1S4GA38 | http://sgid.popgenetics.net/ | *Anopheles gambiae* |
| GLURIIc | A0A1S4G9X1 | http://sgid.popgenetics.net/ | *Anopheles gambiae* |
| 1273165 | F5HKL2 | https://www.uniprot.org/ | *Anopheles gambiae* |
| 1270566 | Q7QHV3 | https://www.uniprot.org/ | *Anopheles gambiae* |
| 1276693 | Q7PNT0 | https://www.uniprot.org/ | *Anopheles gambiae* |
| 1272638 | Q7QDT5 | https://www.uniprot.org/ | *Anopheles gambiae* |

| | | | |
|---|---|---|---|
| 1279695 | Q7PMF1 | https://www.uniprot.org/ | *Anopheles gambiae* |
| GluR4 | AAP41206.1 | https://www.ncbi.nlm.nih.gov/protein/ | *Aplysia californica* |
| GluR3 | NP_001191398.1 | https://www.ncbi.nlm.nih.gov/protein/ | *Aplysia californica* |
| GluR2 | NP_001191540.1 | https://www.ncbi.nlm.nih.gov/protein/ | *Aplysia californica* |
| GluR8 | ACV91074.1 | https://www.ncbi.nlm.nih.gov/protein/ | *Aplysia californica* |
| GluR5 | NP_001191541.1 | https://www.ncbi.nlm.nih.gov/protein/ | *Aplysia californica* |
| GluR1 | NP_001191539.1 | https://www.ncbi.nlm.nih.gov/protein/ | *Aplysia californica* |
| GluR7 | NP_001191543.1 | https://www.ncbi.nlm.nih.gov/protein/ | *Aplysia californica* |
| GluR10 | ACV91076.1 | https://www.ncbi.nlm.nih.gov/protein/ | *Aplysia californica* |
| GluR9 | ACV91075.1 | https://www.ncbi.nlm.nih.gov/protein/ | *Aplysia californica* |
| GluR6 | NP_001191542.1 | https://www.ncbi.nlm.nih.gov/protein/ | *Aplysia californica* |
| GLR1.1 | OAP03138.1 | https://www.ncbi.nlm.nih.gov/protein/ | *Arabidopsis thaliana* |
| GLR1.2 | OAO89985.1 | https://www.ncbi.nlm.nih.gov/protein/ | *Arabidopsis thaliana* |
| GLR1.3 | OAO90809.1 | https://www.ncbi.nlm.nih.gov/protein/ | *Arabidopsis thaliana* |
| GLR1.4 | NP_187408.2 | https://www.ncbi.nlm.nih.gov/protein/ | *Arabidopsis thaliana* |
| GLR2.1 | NP_198062.2 | https://www.ncbi.nlm.nih.gov/protein/ | *Arabidopsis thaliana* |
| GLR2.2 | NP_180048.1 | https://www.ncbi.nlm.nih.gov/protein/ | *Arabidopsis thaliana* |
| GLR2.3 | OAP09775.1 | https://www.ncbi.nlm.nih.gov/protein/ | *Arabidopsis thaliana* |
| GLR2.4 | NP_001320108.1 | https://www.ncbi.nlm.nih.gov/protein/ | *Arabidopsis thaliana* |
| GLR2.5 | OAO89801.1 | https://www.ncbi.nlm.nih.gov/protein/ | *Arabidopsis thaliana* |
| GLR2.6 | NP_001330285.1 | https://www.ncbi.nlm.nih.gov/protein/ | *Arabidopsis thaliana* |
| GLR2.7 | OAP11367.1 | https://www.ncbi.nlm.nih.gov/protein/ | *Arabidopsis thaliana* |
| GLR2.8 | OAP09651.1 | https://www.ncbi.nlm.nih.gov/protein/ | *Arabidopsis thaliana* |
| GLR2.9 | OAP07911.1 | https://www.ncbi.nlm.nih.gov/protein/ | *Arabidopsis thaliana* |
| GLR3.1 | OAP10589.1 | https://www.ncbi.nlm.nih.gov/protein/ | *Arabidopsis thaliana* |
| GLR3.2 | NP_001320141.1 | https://www.ncbi.nlm.nih.gov/protein/ | *Arabidopsis thaliana* |
| GLR3.3 | Q9C8E7 | https://www.uniprot.org/ | *Arabidopsis thaliana* |
| GLR3.4 | NP_001030971.1 | https://www.ncbi.nlm.nih.gov/protein/ | *Arabidopsis thaliana* |
| GLR3.5 | NP_565743.1 | https://www.ncbi.nlm.nih.gov/protein/ | *Arabidopsis thaliana* |
| GLR3.6 | AEE78797.1 | https://www.ncbi.nlm.nih.gov/protein/ | *Arabidopsis thaliana* |
| NMR-1 | AAK01101.2 | https://www.ncbi.nlm.nih.gov/protein/ | *Caenorhabditis elegans* |
| NMR-2 | AAK01102.2 | https://www.ncbi.nlm.nih.gov/protein/ | *Caenorhabditis elegans* |
| GLR-1 | CCD62564.1 | https://www.ncbi.nlm.nih.gov/protein/ | *Caenorhabditis elegans* |
| GLR-2 | AAK01094.2 | https://www.ncbi.nlm.nih.gov/protein/ | *Caenorhabditis elegans* |
| GLR-3 | CAA99883.3 | https://www.ncbi.nlm.nih.gov/protein/ | *Caenorhabditis elegans* |
| GLR-4 | CCD61468.1 | https://www.ncbi.nlm.nih.gov/protein/ | *Caenorhabditis elegans* |
| GLR-5 | CCD62097.1 | https://www.ncbi.nlm.nih.gov/protein/ | *Caenorhabditis elegans* |
| GLR-6 | CCD67907.1 | https://www.ncbi.nlm.nih.gov/protein/ | *Caenorhabditis elegans* |
| GLR-7 | SKC30519.1 | https://www.ncbi.nlm.nih.gov/protein/ | *Caenorhabditis elegans* |

| | | | |
|---|---|---|---|
| GLR-8 | VAY52588.1 | https://www.ncbi.nlm.nih.gov/protein/ | *Caenorhabditis elegans* |
| GRIN1a | XP_021324586.1 | https://www.ncbi.nlm.nih.gov/protein/ | *Danio rerio* |
| GRIN1b | XP_005171833.1 | https://www.ncbi.nlm.nih.gov/protein/ | *Danio rerio* |
| GRIN2A | XP_021329529.1 | https://www.ncbi.nlm.nih.gov/protein/ | *Danio rerio* |
| GRIN2B | XP_017210497.1 | https://www.ncbi.nlm.nih.gov/protein/ | *Danio rerio* |
| GRIN2C | NP_001352716.1 | https://www.ncbi.nlm.nih.gov/protein/ | *Danio rerio* |
| GRIN2D | XP_009292354.1 | https://www.ncbi.nlm.nih.gov/protein/ | *Danio rerio* |
| GRIA1 | AAI62505.1 | https://www.ncbi.nlm.nih.gov/protein/ | *Danio rerio* |
| GRIA2 | XP_005170953.1 | https://www.ncbi.nlm.nih.gov/protein/ | *Danio rerio* |
| GRIA3 | XP_005157190.1 | https://www.ncbi.nlm.nih.gov/protein/ | *Danio rerio* |
| GRIA4 | XP_017214716.1 | https://www.ncbi.nlm.nih.gov/protein/ | *Danio rerio* |
| GRIK1 | XP_009303594.1 | https://www.ncbi.nlm.nih.gov/protein/ | *Danio rerio* |
| GRIK2 | XP_021322472.1 | https://www.ncbi.nlm.nih.gov/protein/ | *Danio rerio* |
| GRIK3 | XP_017206939.1 | https://www.ncbi.nlm.nih.gov/protein/ | *Danio rerio* |
| GRIK4 | E7F1P4 | https://www.uniprot.org/ | *Danio rerio* |
| GRIK5 | A0A0R4IG83 | https://www.uniprot.org/ | *Danio rerio* |
| GRID1 | XP_021323097.1 | https://www.ncbi.nlm.nih.gov/protein/ | *Danio rerio* |
| GRID2 | XP_005167218.1 | https://www.ncbi.nlm.nih.gov/protein/ | *Danio rerio* |
| NMDAR1 | NP_730940.1 | https://www.ncbi.nlm.nih.gov/protein/ | *Drosophila melanogaster* |
| NMDAR2 | NP_001014714.1 | https://www.ncbi.nlm.nih.gov/protein/ | *Drosophila melanogaster* |
| GLURIIA | NP_523484.2 | https://www.ncbi.nlm.nih.gov/protein/ | *Drosophila melanogaster* |
| GLURIIB | AAF52269.3 | https://www.ncbi.nlm.nih.gov/protein/ | *Drosophila melanogaster* |
| GLURIIC | NP_608557.4 | https://www.ncbi.nlm.nih.gov/protein/ | *Drosophila melanogaster* |
| GLURIID | AAG22164.2 | https://www.ncbi.nlm.nih.gov/protein/ | *Drosophila melanogaster* |
| GLURIIE | ABI31184.1 | https://www.ncbi.nlm.nih.gov/protein/ | *Drosophila melanogaster* |
| CG11155 | AAN06582.3 | https://www.ncbi.nlm.nih.gov/protein/ | *Drosophila melanogaster* |
| CG3822 | NP_650925.1 | https://www.ncbi.nlm.nih.gov/protein/ | *Drosophila melanogaster* |
| GRIK | ABI31186.2 | https://www.ncbi.nlm.nih.gov/protein/ | *Drosophila melanogaster* |
| EKAR | ABI29182.1 | https://www.ncbi.nlm.nih.gov/protein/ | *Drosophila melanogaster* |
| Clumsy | ABI31327.1 | https://www.ncbi.nlm.nih.gov/protein/ | *Drosophila melanogaster* |
| GLURIA | Q03445 | https://www.uniprot.org/ | *Drosophila melanogaster* |
| GluRIB | Q9VSV5 | https://www.uniprot.org/ | *Drosophila melanogaster* |
| Ir8a | Q9W365 | https://www.uniprot.org/ | *Drosophila melanogaster* |
| Ir25a | E9NA96 | https://www.uniprot.org/ | *Drosophila melanogaster* |
| GRIN1 | T2ME34 | https://www.uniprot.org/ | *Hydra vulgaris* |

| | | | |
|---|---|---|---|
| GRIN1 | BAI22780.1 | https://www.ncbi.nlm.nih.gov/protein/ | *Pan troglodytes* |
| GRIN2A | XP_016783987.1 | https://www.ncbi.nlm.nih.gov/protein/ | *Pan troglodytes* |
| GRIN2B | XP_016778489.1 | https://www.ncbi.nlm.nih.gov/protein/ | *Pan troglodytes* |
| GRIN2C | XP_016788369.2 | https://www.ncbi.nlm.nih.gov/protein/ | *Pan troglodytes* |
| GRIN2D | C7G3M4 | https://www.uniprot.org/ | *Pan troglodytes* |
| GRIA1 | JAA28648.1 | https://www.ncbi.nlm.nih.gov/protein/ | *Pan troglodytes* |
| GRIA2 | PNI63324.1 | https://www.ncbi.nlm.nih.gov/protein/ | *Pan troglodytes* |
| GRIA3 | XP_016798775.1 | https://www.ncbi.nlm.nih.gov/protein/ | *Pan troglodytes* |
| GRIA4 | XP_016777391.1 | https://www.ncbi.nlm.nih.gov/protein/ | *Pan troglodytes* |
| GRIK1 | XP_009422708.1 | https://www.ncbi.nlm.nih.gov/protein/ | *Pan troglodytes* |
| GRIK2 | XP_009449945.1 | https://www.ncbi.nlm.nih.gov/protein/ | *Pan troglodytes* |
| GRIK3 | XP_524666.3 | https://www.ncbi.nlm.nih.gov/protein/ | *Pan troglodytes* |
| GRIK4 | XP_016775334.1 | https://www.ncbi.nlm.nih.gov/protein/ | *Pan troglodytes* |
| GRIK5 | XP_016789714.2 | https://www.ncbi.nlm.nih.gov/protein/ | *Pan troglodytes* |
| GRID1 | BAI22778.1 | https://www.ncbi.nlm.nih.gov/protein/ | *Pan troglodytes* |
| GRID2 | BAI22779.1 | https://www.ncbi.nlm.nih.gov/protein/ | *Pan troglodytes* |
| PHYPA_015858 | PNR43477.1 | https://www.ncbi.nlm.nih.gov/protein/ | *Physcomitrella patens* |
| PHYPA_020228 | PNR39948.1 | https://www.ncbi.nlm.nih.gov/protein/ | *Physcomitrella patens* |
| GRIN1 | Q62648 | https://www.uniprot.org/ | *Rattus norvegicus* |
| GRIN2A | Q00959 | https://www.uniprot.org/ | *Rattus norvegicus* |
| GRIN2B | Q00960 | https://www.uniprot.org/ | *Rattus norvegicus* |
| GRIN2C | Q00961 | https://www.uniprot.org/ | *Rattus norvegicus* |
| GRIN2D | Q62645 | https://www.uniprot.org/ | *Rattus norvegicus* |
| GRIN3A | Q9R1M7 | https://www.uniprot.org/ | *Rattus norvegicus* |
| GRIN3B | Q8VHN2 | https://www.uniprot.org/ | *Rattus norvegicus* |
| GRIA1 | P19490 | https://www.uniprot.org/ | *Rattus norvegicus* |
| GRIA2 | P19491 | https://www.uniprot.org/ | *Rattus norvegicus* |
| GRIA3 | P19492 | https://www.uniprot.org/ | *Rattus norvegicus* |
| GRIA4 | P19493 | https://www.uniprot.org/ | *Rattus norvegicus* |
| GRIK1 | P22756 | https://www.uniprot.org/ | *Rattus norvegicus* |
| GRIK2 | P42260 | https://www.uniprot.org/ | *Rattus norvegicus* |
| GRIK3 | NP_852038.2 | https://www.ncbi.nlm.nih.gov/protein/ | *Rattus norvegicus* |
| GRIK4 | XP_017450944.1 | https://www.ncbi.nlm.nih.gov/protein/ | *Rattus norvegicus* |
| GRIK5 | EDM08047.1 | https://www.ncbi.nlm.nih.gov/protein/ | *Rattus norvegicus* |
| GRID1 | NP_077354.1 | https://www.ncbi.nlm.nih.gov/protein/ | *Rattus norvegicus* |
| GRID2 | XP_017448392.1 | https://www.ncbi.nlm.nih.gov/protein/ | *Rattus norvegicus* |
| GRIN1 | NP_001081615.1 | https://www.ncbi.nlm.nih.gov/protein/ | *Xenopus laevis* |
| GRIN2A | NP_001106367.1 | https://www.ncbi.nlm.nih.gov/protein/ | *Xenopus laevis* |
| GRIN2B | A7XY94.1 | https://www.uniprot.org/ | *Xenopus laevis* |
| GRIA1 | NP_001153151.2 | https://www.ncbi.nlm.nih.gov/protein/ | *Xenopus laevis* |
| GRIA2 | NP_001153152.1 | https://www.ncbi.nlm.nih.gov/protein/ | *Xenopus laevis* |
| GRIA3 | XP_018084050.1 | https://www.ncbi.nlm.nih.gov/protein/ | *Xenopus laevis* |
| GRIA4 | NP_001153157.1 | https://www.ncbi.nlm.nih.gov/protein/ | *Xenopus laevis* |
| GRID1 | XP_018080997.1 | https://www.ncbi.nlm.nih.gov/protein/ | *Xenopus laevis* |
| GRID2 | XP_018107662.1 | https://www.ncbi.nlm.nih.gov/protein/ | *Xenopus laevis* |
| GLR2.8 | PWZ13255.1 | https://www.ncbi.nlm.nih.gov/protein/ | *Zea mays* |

| | | | |
|---|---|---|---|
| GLR2.9 | PWZ39457.1 | https://www.ncbi.nlm.nih.gov/protein/ | *Zea mays* |
| GLR2.7 | ONM22087.1 | https://www.ncbi.nlm.nih.gov/protein/ | *Zea mays* |
| GLR3.4 | AQK66314.1 | https://www.ncbi.nlm.nih.gov/protein/ | *Zea mays* |
| GLR3.3 | AQK67877.1 | https://www.ncbi.nlm.nih.gov/protein/ | *Zea mays* |
| GRIN1 | XP_020907505.1 | https://www.ncbi.nlm.nih.gov/protein/ | *Exaiptasia pallida* |
| GRIN2A | XP_020900633.1 | https://www.ncbi.nlm.nih.gov/protein/ | *Exaiptasia pallida* |
| GRIN2C | KXJ18695.1 | https://www.ncbi.nlm.nih.gov/protein/ | *Exaiptasia pallida* |
| GRIA1 | XP_020908474.1 | https://www.ncbi.nlm.nih.gov/protein/ | *Exaiptasia pallida* |
| GRIA2 | KXJ23459.1 | https://www.ncbi.nlm.nih.gov/protein/ | *Exaiptasia pallida* |
| GRIA4 | KXJ09774.1 | https://www.ncbi.nlm.nih.gov/protein/ | *Exaiptasia pallida* |
| GRIK2 | XP_020892927.1 | https://www.ncbi.nlm.nih.gov/protein/ | *Exaiptasia pallida* |
| GLR3.5 | KXJ21832.1 | https://www.ncbi.nlm.nih.gov/protein/ | *Exaiptasia pallida* |
| GRIE1 | ML03683a | https://metazoa.ensembl.org/index.html | *Mnemiopsis leidyi* |
| GRIE2 | ML027316a | https://metazoa.ensembl.org/index.html | *Mnemiopsis leidyi* |
| GRIE3 | ML06406a | https://metazoa.ensembl.org/index.html | *Mnemiopsis leidyi* |
| GRIE4 | ML150010a | https://metazoa.ensembl.org/index.html | *Mnemiopsis leidyi* |
| GRIE5 | ML150012a | https://metazoa.ensembl.org/index.html | *Mnemiopsis leidyi* |
| GRIE6 | ML032221a | https://metazoa.ensembl.org/index.html | *Mnemiopsis leidyi* |
| GRIE7 | ML032222a | https://metazoa.ensembl.org/index.html | *Mnemiopsis leidyi* |
| GRIE8 | ML00626a | https://metazoa.ensembl.org/index.html | *Mnemiopsis leidyi* |
| GRIE9 | ML085016a | https://metazoa.ensembl.org/index.html | *Mnemiopsis leidyi* |
| GRIE10 | ML085018a | https://metazoa.ensembl.org/index.html | *Mnemiopsis leidyi* |
| GRIE11 | ML15636a | https://metazoa.ensembl.org/index.html | *Mnemiopsis leidyi* |
| GRIE12 | ML111714a | https://metazoa.ensembl.org/index.html | *Mnemiopsis leidyi* |
| GRIE13 | ML05909a | https://metazoa.ensembl.org/index.html | *Mnemiopsis leidyi* |
| GRIE14 | ML30697a | https://metazoa.ensembl.org/index.html | *Mnemiopsis leidyi* |
| GRIE15 | ML00441a | https://metazoa.ensembl.org/index.html | *Mnemiopsis leidyi* |
| GRIE16 | ML01135a | https://metazoa.ensembl.org/index.html | *Mnemiopsis leidyi* |
| GRIE17 | ML141757a | https://metazoa.ensembl.org/index.html | *Mnemiopsis leidyi* |
| GRIE1 | TriadG55165 | https://metazoa.ensembl.org/index.html | *Trichoplax adhaerens* |
| GRIE2 | TriadG3218 | https://metazoa.ensembl.org/index.html | *Trichoplax adhaerens* |
| GRIE3 | TriadG55165 | https://metazoa.ensembl.org/index.html | *Trichoplax adhaerens* |
| GRIE4 | TriadG61396 | https://metazoa.ensembl.org/index.html | *Trichoplax adhaerens* |
| GRIE5 | TriadG30609 | https://metazoa.ensembl.org/index.html | *Trichoplax adhaerens* |
| GRIE6 | TriadG30612 | https://metazoa.ensembl.org/index.html | *Trichoplax adhaerens* |
| GRIE7 | TriadG18823 | https://metazoa.ensembl.org/index.html | *Trichoplax adhaerens* |
| GRIE8 | TriadG18488 | https://metazoa.ensembl.org/index.html | *Trichoplax adhaerens* |
| GRIE9 | TriadG18943 | https://metazoa.ensembl.org/index.html | *Trichoplax adhaerens* |
| GRIAKDF1 | TriadG19383 | https://metazoa.ensembl.org/index.html | *Trichoplax adhaerens* |
| GRIAKDF2 | TriadT14565 | https://metazoa.ensembl.org/index.html | *Trichoplax adhaerens* |
| GRIAKDF3 | TriadG25025 | https://metazoa.ensembl.org/index.html | *Trichoplax adhaerens* |
| GRIAKDF4 | TriadT25027 | https://metazoa.ensembl.org/index.html | *Trichoplax adhaerens* |
| Griakdf1 | aug_v2a.10154 | https://marinegenomics.oist.jp/coral/ | *Acropora digitifera* |
| Griakdf2 | aug_v2a.07671 | https://marinegenomics.oist.jp/coral/ | *Acropora digitifera* |
| Griakdf3 | aug_v2a.07672 | https://marinegenomics.oist.jp/coral/ | *Acropora digitifera* |
| Griakdf4 | aug_v2a.02216 | https://marinegenomics.oist.jp/coral/ | *Acropora digitifera* |

| | | | |
|---|---|---|---|
| Griakdf5 | aug_v2a.20976 | https://marinegenomics.oist.jp/coral/ | *Acropora digitifera* |
| Grie1 | aug_v2a.11422 | https://marinegenomics.oist.jp/coral/ | *Acropora digitifera* |
| Grie2 | aug_v2a.21171 | https://marinegenomics.oist.jp/coral/ | *Acropora digitifera* |
| Grie3 | aug_v2a.21425 | https://marinegenomics.oist.jp/coral/ | *Acropora digitifera* |
| Grie4 | aug_v2a.00972 | https://marinegenomics.oist.jp/coral/ | *Acropora digitifera* |
| Grincn1 | aug_v2a.06015 | https://marinegenomics.oist.jp/coral/ | *Acropora digitifera* |
| Grincn2 | aug_v2a.22940 | https://marinegenomics.oist.jp/coral/ | *Acropora digitifera* |
| Grincn3 | aug_v2a.13573 | https://marinegenomics.oist.jp/coral/ | *Acropora digitifera* |
| Grin1 | aug_v2a.13302 | https://marinegenomics.oist.jp/coral/ | *Acropora digitifera* |
| Griakdf1 | comp40456_c1_seq62:2-2548(+) | http://www.compagen.org/ | *Oscarella carmela* |
| Griakdf2 | comp40502_c0_seq5:23-2515(+) | http://www.compagen.org/ | *Oscarella carmela* |
| Griakdf3 | comp34412_c0_seq1:2-1051(+) | http://www.compagen.org/ | *Oscarella carmela* |
| Griakdf4 | comp40790_c0_seq11:3-2450(+) | http://www.compagen.org/ | *Oscarella carmela* |
| Griakdf5 | comp33732_c0_seq7:2-2713(+) | http://www.compagen.org/ | *Oscarella carmela* |
| Griakdf6 | comp34218_c0_seq7:170-2386(+) | http://www.compagen.org/ | *Oscarella carmela* |
| Griakdf7 | comp40301_c1_seq41:410-3118(+) | http://www.compagen.org/ | *Oscarella carmela* |
| Griakdf8 | comp33476_c0_seq5:58-2646(+) | http://www.compagen.org/ | *Oscarella carmela* |
| Griakdf9 | comp35603_c0_seq2:3-2636(+) | http://www.compagen.org/ | *Oscarella carmela* |
| Griakdf10 | comp32757_c0_seq1:1-1476(+) | http://www.compagen.org/ | *Oscarella carmela* |
| Griakdf11 | comp36546_c0_seq2:3-980(+) | http://www.compagen.org/ | *Oscarella carmela* |
| Griakdf12 | comp39312_c0_seq2:2-1471(+) | http://www.compagen.org/ | *Oscarella carmela* |
| Gril1 | comp20375_c0_seq1:2-2641(+) | http://www.compagen.org/ | *Oscarella carmela* |
| Gril2 | comp32525_c0_seq2:1-2547(+) | http://www.compagen.org/ | *Oscarella carmela* |
| Gril3 | comp37028_c0_seq1:2-2653(+) | http://www.compagen.org/ | *Oscarella carmela* |
| Gril4 | comp39946_c0_seq2:3-2558(+) | http://www.compagen.org/ | *Oscarella carmela* |
| Gril5 | comp40747_c0_seq5:168-2879(+) | http://www.compagen.org/ | *Oscarella carmela* |
| Gril6 | comp7940_c0_seq1:1-2721(+) | http://www.compagen.org/ | *Oscarella carmela* |
| Gril7 | comp42765_c0_seq1:2-2815(+) | http://www.compagen.org/ | *Oscarella carmela* |
| GRIK1 | scpid25255\| scgid10696 | http://www.compagen.org/ | *Sycon cilliatum* |
| GRIL1 | scpid21909\| scgid9412 | http://www.compagen.org/ | *Sycon cilliatum* |
| GRIL2 | scpid27594\| scgid17785 | http://www.compagen.org/ | *Sycon cilliatum* |
| GRIL3 | scpid25297\| scgid5997 | http://www.compagen.org/ | *Sycon cilliatum* |
| GRIL4 | scpid22929\| scgid17334 | http://www.compagen.org/ | *Sycon cilliatum* |

| | | | |
|---|---|---|---|
| GluR0 | P73797 | https://www.uniprot.org/ | *Synechocystis sp. PCC 6803* |
| BGIBMGA010135-RA | BGIBMGA010135-RA | http://sgid.popgenetics.net/ | *Bombyx mori* |
| BGIBMGA011691-RA | BGIBMGA011691-RA | http://sgid.popgenetics.net/ | *Bombyx mori* |
| BGIBMGA011484-RA | BGIBMGA011484-RA | http://sgid.popgenetics.net/ | *Bombyx mori* |
| BGIBMGA008721 | BGIBMGA008722 | http://sgid.popgenetics.net/ | *Bombyx mori* |
| BGIBMGA011590-RA | BGIBMGA011590-RA | http://sgid.popgenetics.net/ | *Bombyx mori* |
| BGIBMGA011260-RA | BGIBMGA011260-RA | http://sgid.popgenetics.net/ | *Bombyx mori* |
| BGIBMGA010960-RA | BGIBMGA010960-RA | http://sgid.popgenetics.net/ | *Bombyx mori* |
| BGIBMGA000645-RA | BGIBMGA000645-RA | http://sgid.popgenetics.net/ | *Bombyx mori* |
| BGIBMGA011528-RA | BGIBMGA011528-RA | http://sgid.popgenetics.net/ | *Bombyx mori* |
| GluR2 | B3G464 | https://www.uniprot.org/ | *Adineta vaga* |
| GluR1 | E9P5T5 | https://www.uniprot.org/ | *Adineta vaga* |
